# The role of artificial photo backgrounds of shelter dogs on pet profile clicking and the perception of sociability

**DOI:** 10.1101/2021.08.04.455112

**Authors:** F Lamb, A Andrukonis, A Protopopova

**Author notes:** Corresponding author (AP). These authors contributed equally to the study conception, design, and statistical approaches. All authors reviewed the manuscript. This author wrote the main manuscript text and completed the data acquisition and formal analysis (F Lamb).

## Abstract

With the increasing prevalence of technology, the internet is often the first step for potential pet owners searching for an adoptable dog. However, best practices for the online portrayal of shelter and foster dogs remain unclear. Different online photo backgrounds appearing on adoption websites for shelter dogs may impact adoption speed by influencing viewer interest. Online clicking behaviour on pet profiles and human-directed sociability, broadly defined, has been previously linked to increased adoption likelihood. Therefore, the objective of this study was to determine the relationship between photo backgrounds of shelter dogs and online clicking as well as perceived human-directed sociability. In a virtual experiment, 680 participants were asked to rank the sociability and friendliness of four different adoptable dogs on a scale from 0-10. The photo background of each dog was digitally altered and randomly assigned to four experimental background conditions: 1) outdoor, 2) home indoor, 3) in-kennel indoor, and 4) plain coloured. As a proxy for adoption interest, a link to the dog’s adoption profile was presented on each slide and the clicking behaviour of participants on this link was recorded. Mixed logistic regression and poisson models revealed that background did not affect participants’ link-clicking behaviour (*chisq = 3*.*55, df = 3, p = .314*) nor perceptions of sociability (*statistic = 6*.*19, df = 3, p = .103*). Across all backgrounds, only 4.74% of presented slides culminated in participant link-clicking. Sociability scores also did not predict link clicking. Assessment of participant-related factors and dog ID revealed that link-clicking and sociability scores of photographs were influenced by differences between dogs themselves and unaffected by participants’ awareness of study hypotheses. We conclude that artificial background types did not affect participant responses. The results demonstrate the importance of empirical data in making marketing decisions in animal shelters. Understanding which aspects of online marketing materials impact viewer interest will provide guidance for both animal shelter personnel and foster families to improve speed of adoption of the animals in their care.

## Introduction

Between 3.9-5.5 million dogs in the United States, 28,000 dogs in Canada, and unknown numbers across the world, enter animal rehoming shelters annually [1–3]. Dogs within shelters are subjected to non-ideal conditions: an unfamiliar environment, minimal social interactions, noise disturbances, and many other potential stressors [4–6]. Given that the environmental stressors within animal shelters may be difficult or impossible to mitigate, many shelter professionals now advocate focusing programs to reduce the length of stay of animals sheltered. With technology now a common feature integrated into everyday life [7,8], and likewise integrated into animal sheltering, shelters can expand their reach to potential pet adopters online and reduce the time until adoption for the animals in their care. Online platforms such as Petfinder or social media sites advertise adoptable animals to the public [9]. Pet owners consider these internet adoption platforms as valuable resources; 36% of dog adopters and 30% of cat adopters reported that access to information from Petfinder or shelter websites provided important sources of information about their pet prior to adoption [10]. Optimizing these online adoption platforms could be a useful tool to reduce the length of stay for animals in shelters.

However, it is important to consider the type of online advertisements for shelter animals as certain factors can increase or decrease the interest of potential adopters, thereby influencing the likelihood of adoptions [9,11–13]. A range of mutable photo traits such as improved photo quality, increased direct eye contact with the camera, presence of a collar, or other accessories such as bandanas, were found to decrease the median days to adoption of shelter dogs [11,13]. Immutable photo traits of shelter dogs such as morphological features relating to coat colour and ear type (floppy or erect) can also impact their length of stay within shelters [12]. Strikingly, the effect of photo traits of companion shelter animals can be measured from the online behaviour of viewers [9]. Past research showed that increased online clicking on a shelter cat profile with different coat colours predicted shorter length of stays for the same cats [9]. Cats with cream-coloured coats had the shortest length of stay and received the highest number of online profile-clicking, whereas cats with black-coloured coats had the longest length of stay and received the lowest number of online profile-clicking [9].

Immutable dog features displayed in online photos have also been implicated in affecting the determination of dog personality traits by potential adopters [14]. For example, floppy-eared dogs were rated high in Agreeableness (e.g. warm) and Emotional Stability (e.g. calm) compared to pointy-eared dogs, whereas dogs with yellow coats were ranked with higher Agreeableness, Conscientiousness (e.g. dependable), and Emotional Stability compared to dogs with black coats [14]. As sociability has been similarly broadly defined as approachable, friendly, intelligent, and less aggressive [15], it is likely that potential adopters can develop a sense of the human-directed sociability of dogs based on photographs within an online environment. With the perceived sociability of dogs previously linked to their likelihood of selection and their length of stay [16,17], sociability scores may be an appropriate proxy for adopter desirability when assessing photographs of shelter dogs. Although immutable photo characteristics, such as the physical appearance of dogs, are often cited as a predictive factor taken into consideration when adopting [10], further attention on mutable photos traits, such as online photo backgrounds, is warranted as these photo elements are modifiable by photographers.

Although previous research has assessed the role of different online photo backgrounds on the speed of adoption, the preferred background type remains unclear [11–13]. While some studies reported that shelter dogs photographed outside of kennels and in outdoor environments increased adoption rates [11,13], others reported the opposite finding, where dogs photographed in natural environments had the longest length of stay at the shelter compared to dogs photographed in a kennel or indoor environment [12]. It is also unclear how plain coloured backgrounds of online photos of shelter dogs impact the speed of adoption. Research is currently lacking in comparing pure coloured backgrounds to outdoor, home indoor, and in-kennel background types. The usage of coloured backgrounds is presently prevalent on pet adoption sites, such as Petfinder, when displaying online photos of shelter dogs. Third-party companies have also emerged claiming that coloured backgrounds can serve to improve adoption rates and increase online engagement. The effect of colour on the online purchasing intention and attitudes of consumers have been previously well investigated in marketing and advertising literature [18]. Viswanathan and Swaminathan found that high colour impact displays on the landing page of web pages can increase click-through-rates [19]. However, it is unclear whether the use of coloured backgrounds may negatively impact the speed of adoption of shelter dogs as previous research has suggested that potential adopters may utilize the background of photographed shelter dogs to infer information influencing their decision to adopt [11,12]. Coloured backgrounds portray dogs within a highly artificial environment, whereby information such as the suitability of shelter dogs within a home environment as assessed from dogs displayed with home indoor backgrounds may not be visually available for online users. As such, further investigation of coloured backgrounds in the context of online advertisement for shelter dogs is warranted.

The objective of the present study was to assess if the background type of online photos of shelter dogs alter proxies of adoption interest: online dog profile clicking behaviour [9] and perceived human-directed sociability (hereafter: sociability scores [15]). A within-subject online experiment was conducted on four background conditions (outdoor, home indoor, in-kennel, plain coloured) to test our primary hypothesis that background type will predict differential clicking on pet profiles and our secondary hypothesis that background type will alter perceptions of sociability. Finally, we examined how participant-related demographic factors (participants’ location of residence; participants’ age category; previous or current dog ownership), awareness of photo manipulation, and dog ID impacted online clicking response on pet profiles and sociability scores on digitally altered dog photographs. Understanding how mutable photo characteristics of shelter dogs affect online interest of potential adopters’ through these measures can provide further guidance and strategies for shelter personnel, volunteers, and foster families in increasing the speed of adoption of animals in their care.

## Materials and Methods

All experimental procedures were approved by the University of British Columbia Behavioural Research Ethics Board (H20-02584).

### Participants

A total of n = 958 participants were recruited from social media sites (Facebook, Twitter) and online community forums (Reddit) to participate in a virtual experiment administered online using the software Qualtrics, which was active online between November 17th, 2020 to November 30th, 2020. Participants who indicated an interest in dogs were required to be aged 18 years or older to provide written consent to this study. No monetary compensation was given to participants completing the online experiment. Due to 270 incomplete responses and eight flagged as potential bot activity (recaptcha scores ≤ 0.5), a total of 680 participants were included in the final analyses. The majority of participants were female (85%) and either current or past dog owners (89.4%). While the participant age category of 25-34 was the highest (23.01%), all age categories were evenly represented in this study; demographics are further described in Table 1.

**Table 1.**
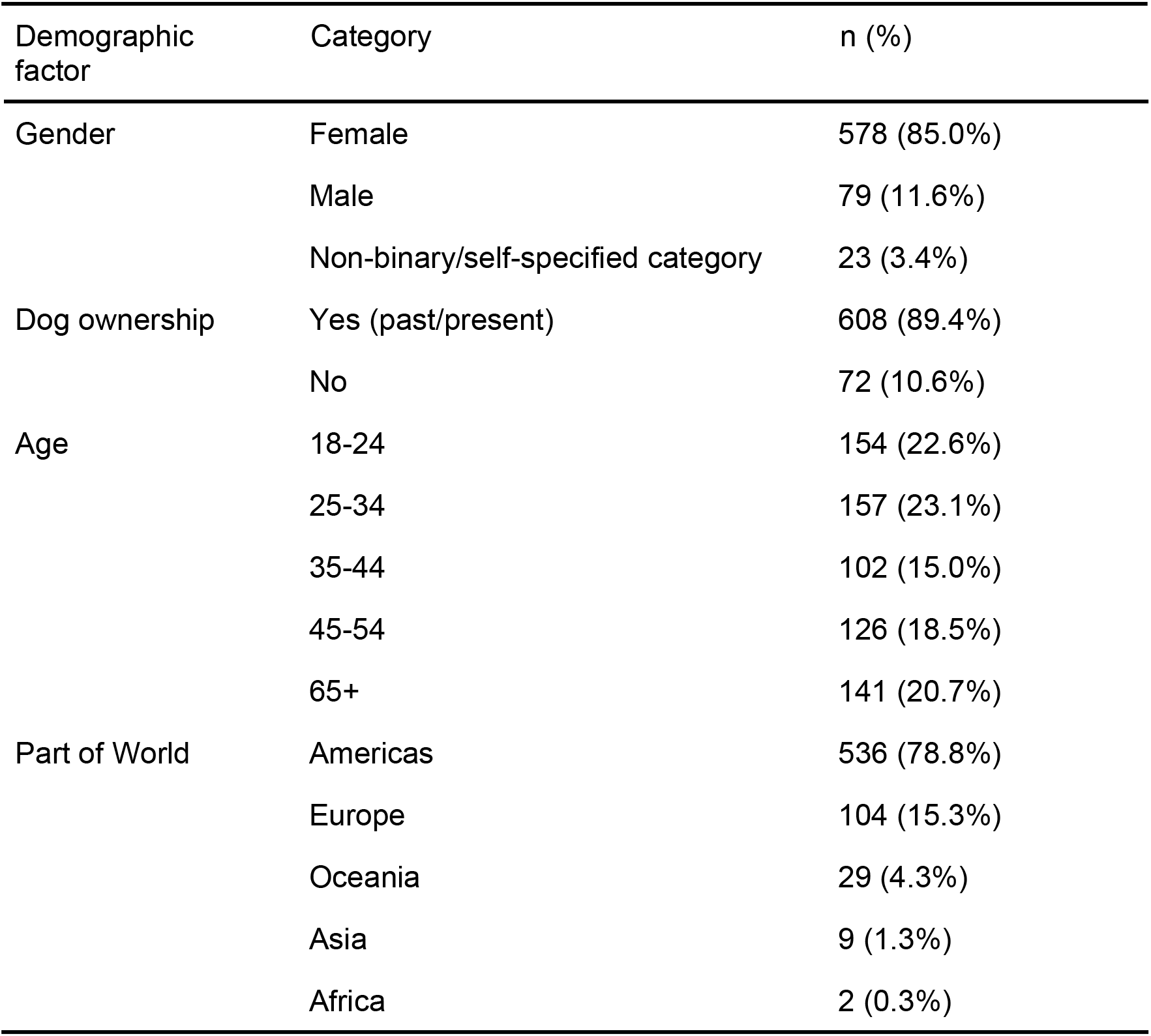
Demographic factors of participants (n = 680)

### Procedure

A total of 680 participants viewed four online photos of adoptable dogs that were selected from the Haven Animal Care Shelter in Lubbock, Texas on Petfinder and digitally altered with Word Office 365. As the majority of dogs entering shelters are mixed breed [29,30], all four photographs were selected as mixed breed dogs. For each dog featured online, there was a 25% chance of the dog being displayed situated in one of the four different conditions: a) outdoor, b) home indoor, c) in-kennel, d) plain coloured. As a result, some participants viewed certain background types more than once. The colour “blue” was selected as the plain coloured background as this colour was preferred in online settings [31] and positively associated with consumer trust [32–34]. Using this within-subject design, individual differences between dogs were taken into account in determining the effect of artificial backgrounds types. All backgrounds were also used consistently across all dogs and the online interface, Qualtrics, permitted participants to view each dog presented in the same order (Liberty; Phantom; Anakin; Rogue) a single time (Fig 1). Participants were initially informed four adoptable dogs would be displayed and were not provided the option to revisit previous questions on Qualtrics. To ensure standardization, dog photographs were selected based on an exclusion criterion designed to maintain consistency among all online photos of dogs selected. This criterion was based on mutable photo traits previously found to impact the speed of adoption or the perception of adoptability/friendliness (Table 2). For consistency among the four dog photographs, accessories such as toys, collars, and bandanas [12,13] and additional photo elements such as the presence of handlers were excluded [35]. All photographs were taken by a professional photographer from the Haven Animal Care Shelter with dogs in a sitting position [24], having direct eye contact with the camera [11], with mouths open [12], of good photo quality (non-blurry; [11], and adequately sized [11]).

**Table 2.** Exclusion criteria for mutable photo traits of dogs.

| Photo trait | Effect on adoption | Exclusion criteria |
| --- | --- | --- |
| Toys | Toy present: decreased speed of adoption [12] | No toys |
| Collar | Collar present: increased speed of adoption [13] | No collars |
| Bandana | Bandana present: decreased speed of adoption [12] | No bandanas |
| Photo quality | Focused: increased speed of adoption [11] | No blurry photos |
| Handler | Elderly women/male child: increased friendliness/adoptability [35] | No handlers |
| Pose | Sitting alone: rated highest for adoptability/friendliness [24] | No dogs standing |
| Mouth | Mouth closed: increase in adoption speed [12] | No dogs with mouth closed |
| Photo size | Small photos: decreased speed of adoption [11] | No small photos |
| Eye contact | Eye contact: increased speed of adoption [11] | No photos with no eye contact with camera |

**Fig 1.**
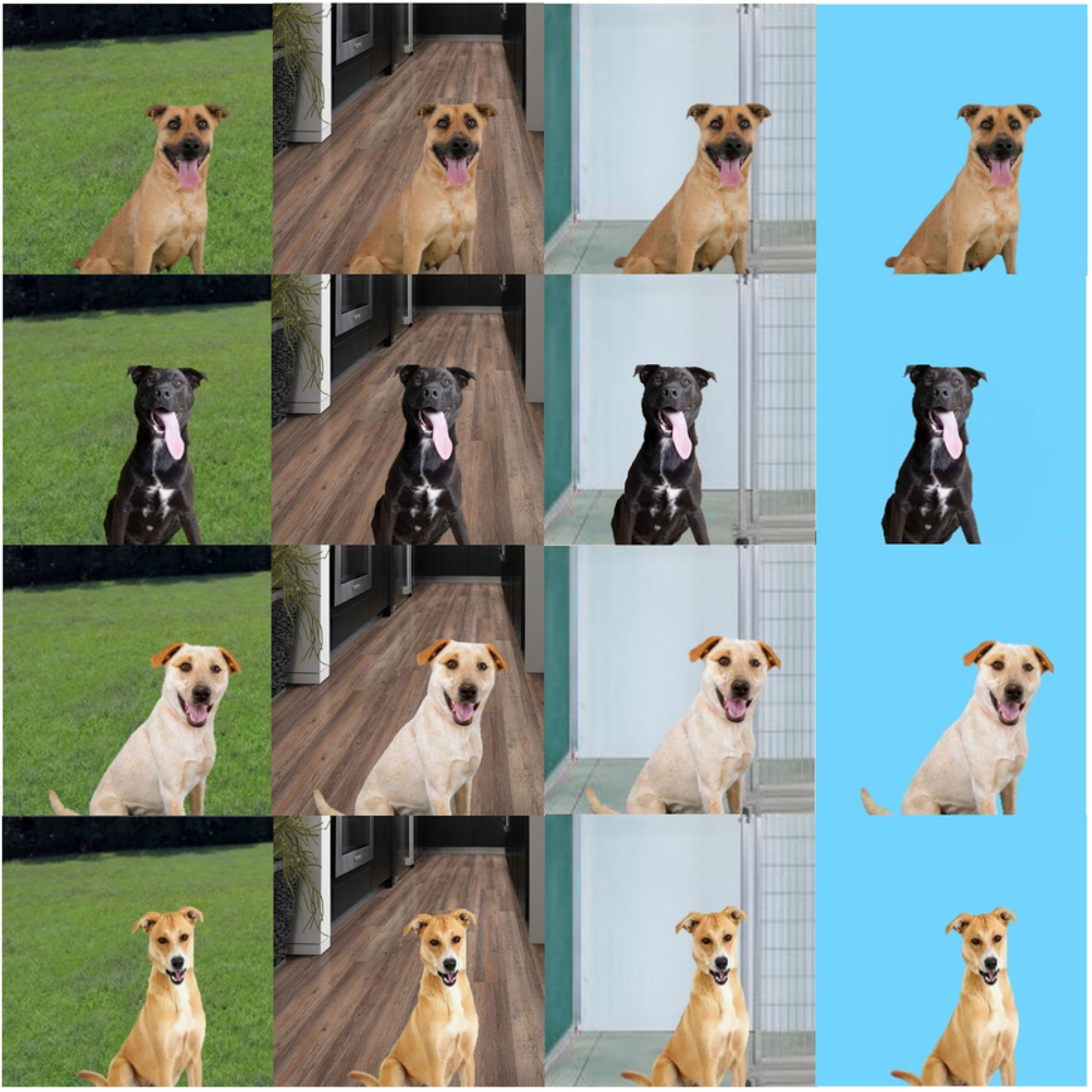
Dogs and backgrounds displayed online. From top to bottom dogs present in this study were: Liberty, Phantom, Anakin, and Rogue. From left to right background types present in this study were: outdoor, home indoor, in-kennel, and plain coloured backgrounds.

For each dog, displayed randomly with either an outdoor, home indoor, in-kennel, or plain coloured background, participants were asked to rank the perceived friendliness/sociability of dogs shown in online photos using a slider on a discrete scale of zero (lowest sociability/friendliness) to ten (highest sociability/friendliness; Fig 2). Participants were also provided a link below each dog photo and informed to access the link if they would like to visit the adoption website of the dog featured. Qualtrics was programmed to record when participants access the link to quantitatively measure adoption interest via link clicking behaviour [9] (Fig 2). However, participants were not informed that online photo backgrounds of each dog were altered nor were they informed that link clicking behaviour was recorded. This information was withheld to prevent participants’ knowledge of online photo backgrounds from influencing their response to the different photos. Instead, participants were initially told that the purpose of the study was to collect data on the human perception of personality of dogs based on online photos to help design an algorithm for a software that could recognize positively perceived dog traits in online photos. Demographic questions were included at the start of the online experiment. For the last question, participant awareness of photo backgrounds being manipulated and link clicking being recorded were measured on a score of 1 (definitely not aware) to 5 (definitely aware) after informing participants of our true study goals.

**Fig 2.**
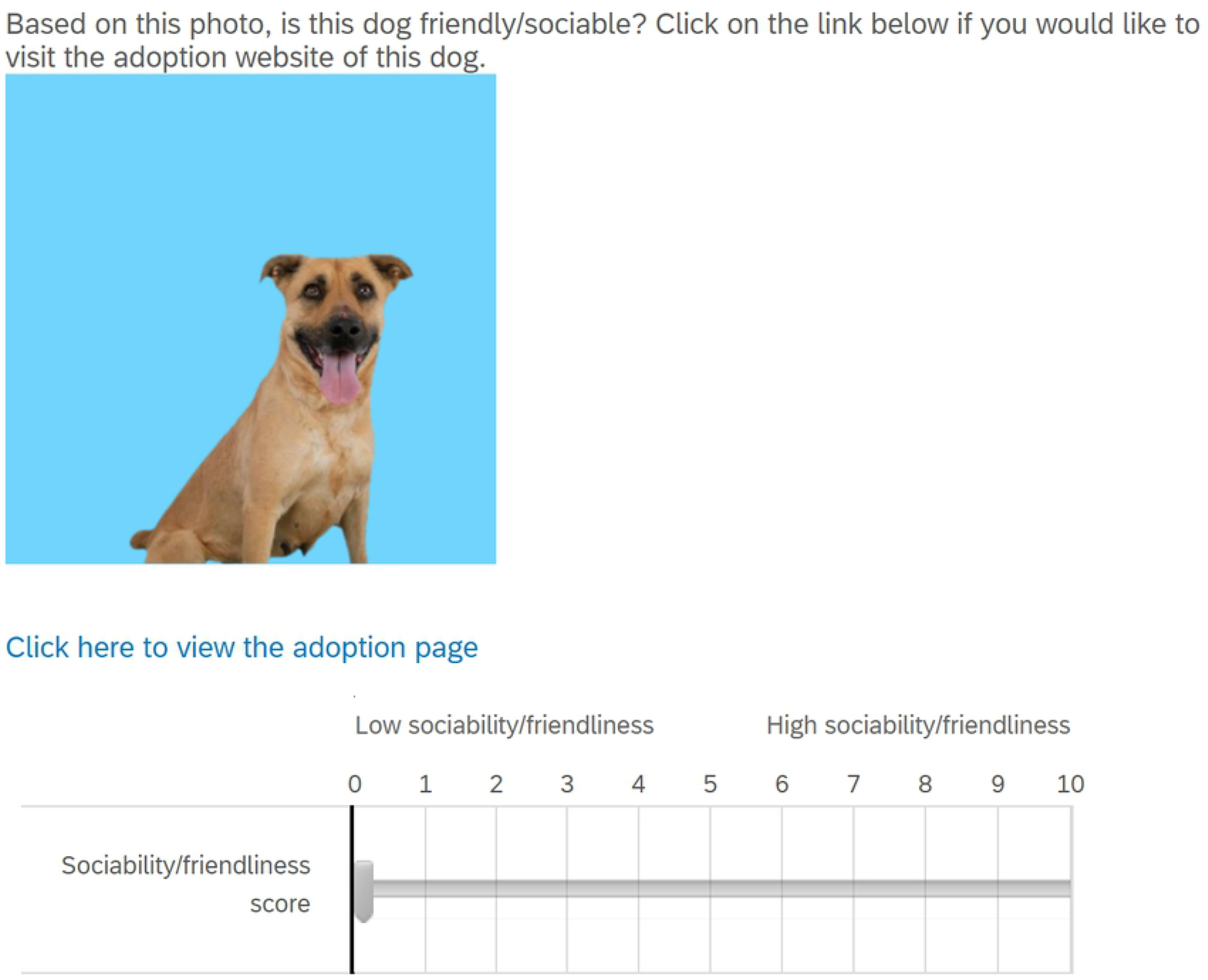
Sample question provided to participants on Qualtrics. For each dog a) a sociability/friendliness score slider is provided and b) a link for participants to access the adoption website specific to the dog featured in the photograph is provided.

### Statistical Analysis

All cleaning of data and statistical analyses were performed using R on n = 2,720 data points [36]. Shapiro-Wilk tests were performed to determine the normality of the sociability scores. For link clicking behaviour, the effect of background, sociability scores (0-10), dog ownership (yes; no), dog ID (Anakin; Rogue; Phantom; Liberty), participant awareness of photo manipulation scores (1-5), participant location of residence (Africa; Americas; Asia; Europe; Oceania), and participant age category (18-24; 25-34; 35-44; 45-54; 65+) were tested using a mixed effects logistic model where participant identity was specified as a random effect. For sociability scores, the effect of background type was tested with a second mixed effects model fitted to a poisson distribution. Background type, dog ownership, dog ID, participant photo manipulation awareness, participant location of residence, participant age category were fixed effects while participant identity was specified as a random effect. Statistical significance was noted if *p <* .05.

## Results

### Link clicking

Type II Wald Chi-Squared Tests on a mixed effects logistic regression model revealed that the background type as a predictor variable of online link clicking behaviour on dog profiles was not statistically significant (*chisq* = 3.55, *df* = 3, *p* = .31). Table 3 shows the number and percentage of slides clicked by condition. In total, 4.74% of presented slides resulted in participant pet profile clicks. There was no effect of sociability scores (*chisq = 4*.*79, df = 10, p = .91*), participants’ location of residence (*chisq = .50, df = 4, p = .97)*, prior or current dog ownership experience (*chisq = .20, df = 1, p = .34*), and participants’ age (*chisq = 3*.*14, df = 4, p = .71*). Participant awareness of photo manipulation scores (*chisq = 1*.*09, df = 4, p = .90*) did not predict link clicking on pet profiles as well (Table 4). Further analysis with a Kruskal-Wallis test revealed that the mean awareness of photo manipulation scores did not differ between background types (*statistic = 6*.*74, df = 3, p = .08*). However, there was a significant relationship between dog ID’ and link clicking behaviour (*chisq = 13*.*78, df = 3, p = .003*). Post Hoc pairwise comparisons revealed that participant link clicking differed significantly between Liberty and Rogue (*p = .025*) and between Anakin and Liberty (*p = .0053*). Specifically, Liberty received the most clicks (6.91%) while Phantom, Rogue, and Anakin received 4.85%, 3.68%, and 3.53% clicks respectively (Fig 3).

**Table 3.**
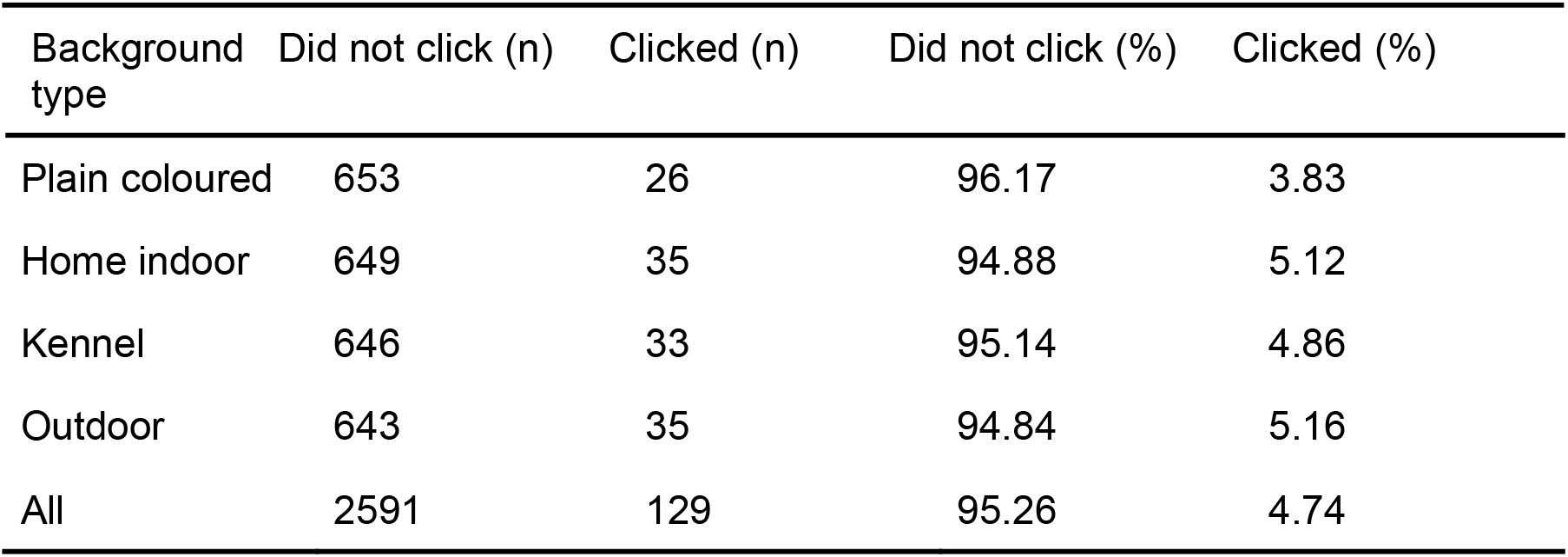
Summary of participant link clicking between background types.

| Background type | Did not click (n) | Clicked (n) | Did not click (%) | Clicked (%) |
| --- | --- | --- | --- | --- |
| Plain coloured | 653 | 26 | 96.17 | 3.83 |
| Home indoor | 649 | 35 | 94.88 | 5.12 |
| Kennel | 646 | 33 | 95.14 | 4.86 |
| Outdoor | 643 | 35 | 94.84 | 5.16 |
| All | 2591 | 129 | 95.26 | 4.74 |

**Table 4.** The number (n) and percentage (%) of participants clicking on pet profiles by photo manipulation awareness scores (1-5).

| Photo manipulation awareness score | Total (n) | Clicked | Clicked (%) |
| --- | --- | --- | --- |
| 1 (Definitely not) | 524 | 25 | 4.77 |
| 2 (Probably not) | 584 | 21 | 3.60 |
| 3 (Might or might not) | 416 | 25 | 6.01 |
| 4 (Probably yes) | 616 | 18 | 2.92 |
| 5 (Definitely yes) | 580 | 40 | 6.90 |

**Fig 3.**
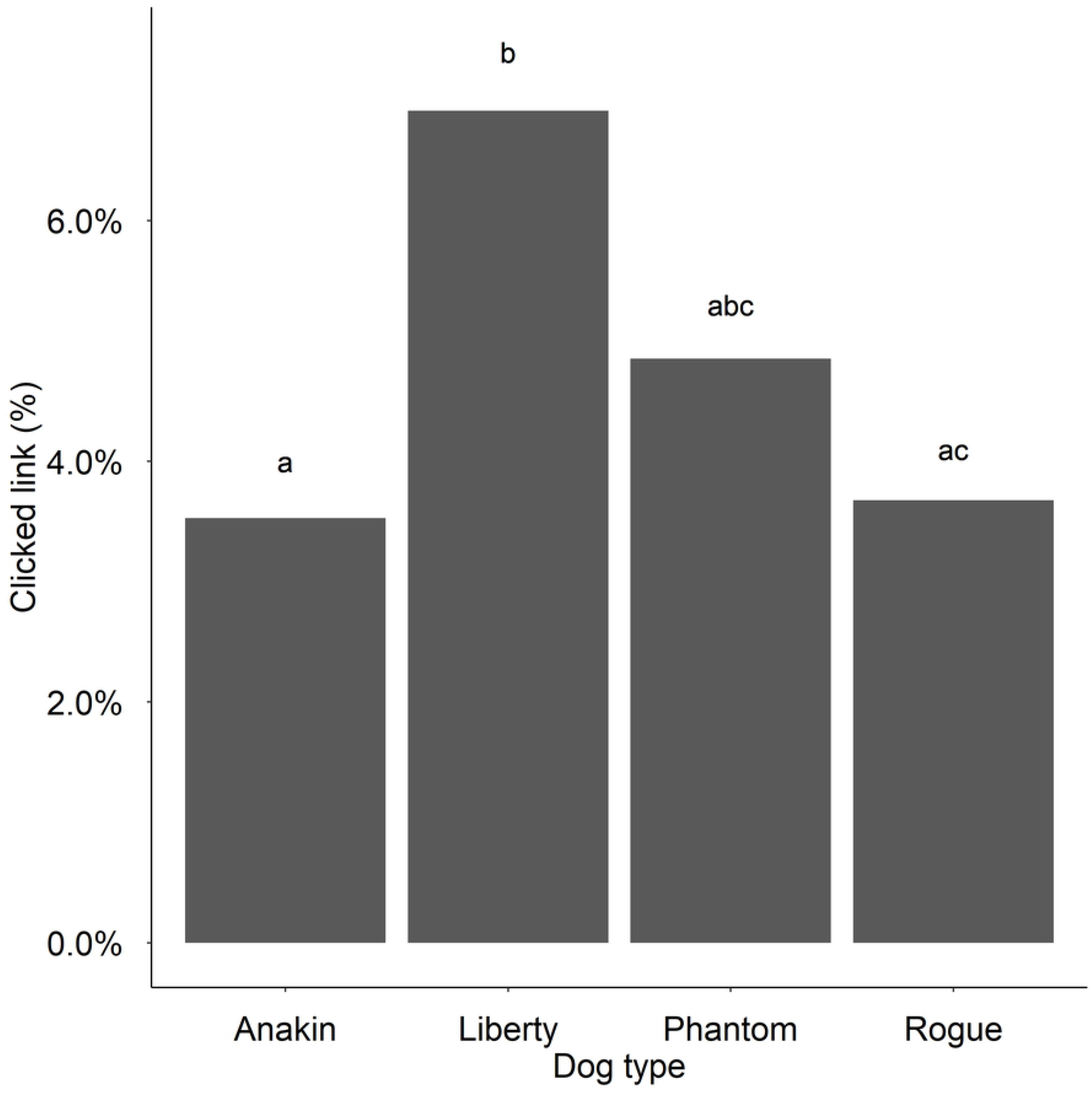
Link clicking by dog type. The percentage of participants clicking on each dog photo presented online.

### Sociability/friendliness score

Type II Wald Chi-Squared Tests on a mixed effects poisson model revealed that background type did not predict sociability scores (*statistic = 2*.*48, df = 3, p = .48*). Specifically, the difference in mean sociability scores for plain coloured (*M = 7*.*00, SD = 1*.*94*), home indoor (*M = 7*.*19, SD = 1*.*91)*, in-kennel (*M = 7*.*02, SD = 2*.*00*), and outdoor (*M = 6*.*98, SD = 1*.*94*) backgrounds were not statistically different. The similar distribution of sociability scores by background type can be visualized in Fig 4. There was no effect of participants’ prior or current dog ownership experience (*chisq = 1*.*13, df = 1, p = .29*), participant age (*chisq = 7*.*81, df = 4, p=*.*099*), and participants’ awareness of photo manipulation scores (*chisq = 0*.*078, df = 4, p = .78*). However, dog ID (*chisq = 81*.*53, df = 3, p = <*.*0001*) and participants’ location of residence (*chisq = 13*.*66, df = 4, p = .0085*) were significant predictors of sociability scores. Mean sociability scores between dog IDs from highest to lowest were the following: Phantom (*M = 7*.*56, SD = 1*.*70*), Liberty (*M = 7*.*23, SD = 2*.*03*), Anakin (*M = 7*.*09, SD = 1*.*94*), Rogue (*M = 6*.*31, SD = 1*.*91*). Post Hoc pairwise comparisons revealed that sociability scores differed between Anakin and Liberty (*p < .0001*), Anakin and Phantom (*p < .0001*), Anakin and Rogue (*p < .0001*), Liberty and Rogue (*p < .0001*), and Phantom and Rogue (*p < .0001*); however, sociability scores did not differ between Liberty and Phantom (*p = .21*). Additional Post Hoc pairwise comparisons for participants’ location of residence revealed that sociability scores differed significantly between Americas (*M = 7*.*16, SD = 1*.*95*) and Europe (*M = 6*.*60, SD = 1*.*91; p = .023*). All other pairwise comparisons between locations (Africa, Europe, Asia, Oceania, Americas) had no significant effect on sociability scores (*p > .05*); see Table 5 for summary values for sociability scores by continent type.

**Table 5.** Sociability scores between continents: sample size (n), mean, standard deviation, median, interquartile range (IQR).

| Continent | n | Mean | Standard deviation | Median | IQR |
| --- | --- | --- | --- | --- | --- |
| Africa | 8 | 6.63 | 2.20 | 6.50 | 4.25 |
| Americas | 2144 | 7.16 | 1.95 | 7.00 | 3.00 |
| Asia | 36 | 7.36 | 1.78 | 8.00 | 2.25 |
| Europe | 416 | 6.60 | 1.91 | 7.00 | 3.00 |
| Oceania | 116 | 6.49 | 1.88 | 7.00 | 3.00 |

**Fig 4.**
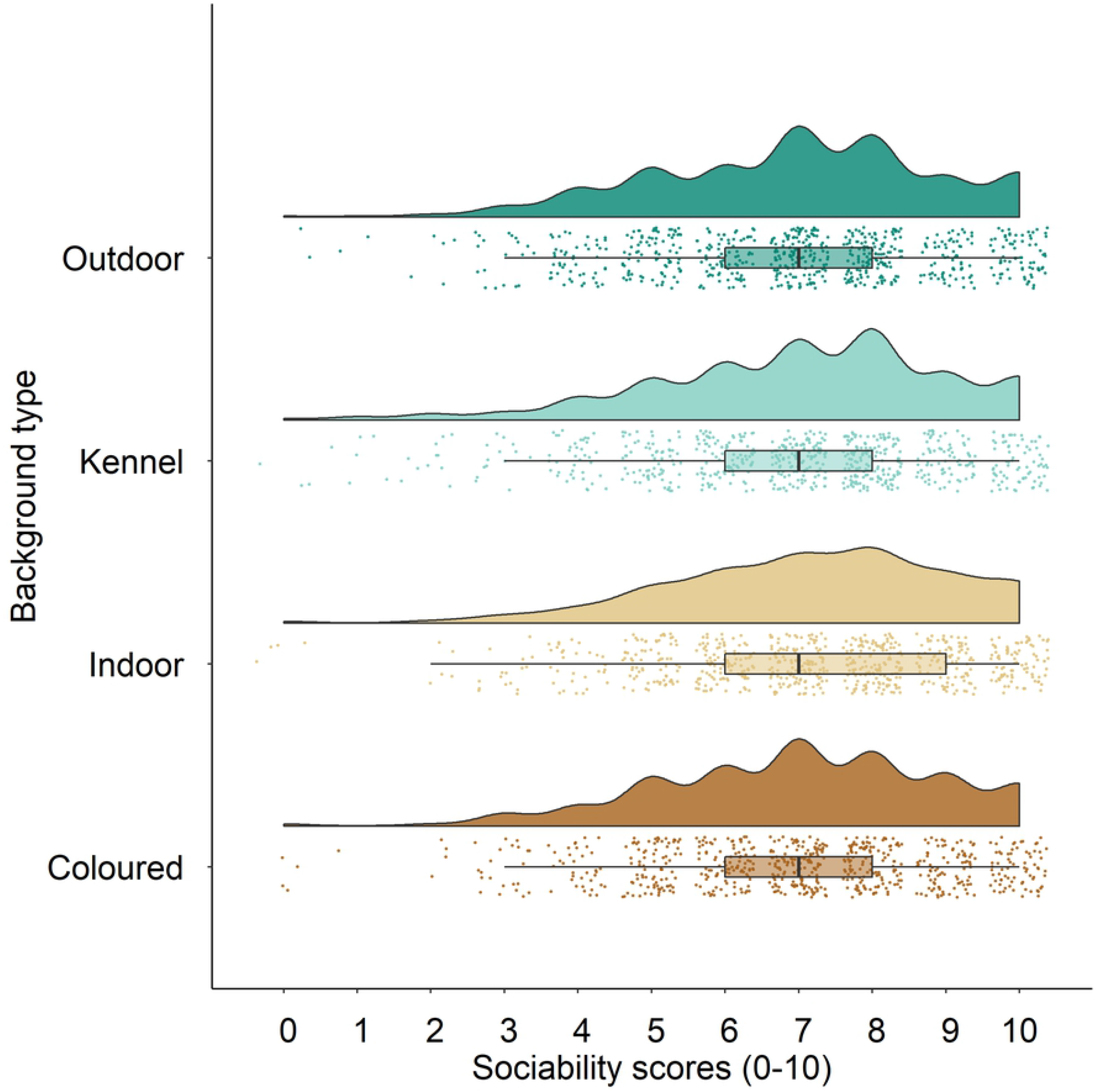
Sociability scores by background type. Perceived sociability scores of photographed dogs by background type visualized with a distribution (shaded portion) and data points (n=2720). The central dot represents the mean while the line across each dot represents the 95% confidence interval.

## Discussion

This study explored the impact of artificial background types of online dog photos on the link clicking engagement with dog profiles and the perceived sociability of shelter dogs among participants with an interest in dogs. It has been previously suggested that background types impact the speed of adoption [11–13]. The results of our study indicated no effect of background type on the initial online engagement with pet profiles nor the initial assessment of sociability by online users; however this is the first online experimental assessment of background types within dogs, to the knowledge of the authors, where background types were digitally altered. Assessment of awareness of photo manipulation in relation to background type, morphological canine attributes, and demographic-related factors in this study provides further guidance for advertising and marketing shelter dogs.

### Background type

Among dogs displayed online in outdoor, home indoor, in-kennel, and plain coloured backgrounds our primary and secondary hypotheses were not supported as background type did not influence link clicking nor sociability scores on dog profiles. The results are in contrast to previous findings where background type in photographs of shelter dogs had an effect on adoption preference or the length of stay of dogs within shelters [11–13]. Nakamura et al. found that shelter dogs photographed in kennel environments had the shortest length of stay [12] while Thorn and Mitchell and Lampe and Witte found that shelter dogs photographed outside of kennel environments had increased adoption interest [13] and decreased length [11] of stay respectively.

Our results may be due to differences of the online interface where participants viewed photographs of shelter dogs and differences of digitally altered photos. Photographs of dogs viewed by participants in this study were provided a single opportunity to view each dog. With 95.26% of links leading to pet profiles not accessed from scored photographs, it is likely that sociability scores were purely based on the assessment of photographs. In other studies, dog photos were taken directly from online adoption directories and differences in background types in relation to the duration of time pet profiles remained active on adoption sites were then analyzed between different dogs in a correlational design [11,12]. Due to the nature of online adoption directories, potential adopters were permitted to view pet profiles multiple times and could view additional information about featured dogs from written descriptions. A recent study reveal<u>ed</u>s that personality adjectives such as “active” or “gentle” within descriptive texts of online profiles for shelter dogs can influence the appeal of certain dog breeds and impact the length of stay [20]. Descriptors relating to friendliness and sociability appeared in 37% of pet profiles [20]. Therefore, it is possible that potential adopters may utilize information from backgrounds of online photos in conjunction with information on pet profiles to formulate perceptions of dogs featured online; however, further research is needed. Alternatively, it is possible that the judgement of participants interested in dogs may not be equivalent to the judgement of participants with the intention of adoption. This may be the case in our study as the criteria for participating in this study was broadly defined and directed to the general public with an interest in dogs. The importance of background type may differ between adopters and non-adopters. While an in-kennel background might appeal to potential adopters intending to adopt to help dogs that “appear to be in need” [12], an in-kennel background may not prompt the same level of emotional response and online interest for non-adopters without these existing motivations for adoption. For future research, perhaps a more targeted group of individuals with the intention of adoption should be included. In this way, the motivation to adopt can be assessed in relation to link clicking and sociability scores ascribed to different artificial backgrounds.

Perceived sociability of dogs in online photos did not predict link clicking on dog profiles. This was unexpected as sociability has previously been described as a desirable trait in dogs that increases the likelihood shelter dogs are selected [16,17]. However, Protopopova et al. found that using passive sociable behaviours such as gazing did not increase adoption rates, whereas using more active sociable behaviours such as lying in proximity to the adopter and engaging in play initiating increased adoption rates [21]. It is possible that online photographs of shelter dogs are well-suited in displaying physical characteristics, but constrained from displaying active behaviours that are more closely linked to sociability [22]. When comparing photographs and videos used to promote the adoption of shelter dogs, dogs displayed with a 30-second video were perceived as more trainable, intelligent, friendly, and gentle as well as less dominant, aggressive, and unsociable [22]. This suggests that the perceived sociability of dogs online may not be a valid proxy for adoption interest specifically for photographs within an online environment. However, future studies should investigate if different backgrounds types will predict differential sociability scoring of shelter dogs displayed in videos.

Along the same vein, photo backgrounds being digitally altered rather than sourced directly from adoption websites may contribute to background type having no effect on sociability scores and link clicking behaviour on pet profiles. For example, Lampe and Witte [11] suggested that outdoor backgrounds may increase the speed of adoption as natural lighting may improve photo quality [11]. With photographs of dogs in this study originally displayed on Petfinder with an indoor background, all photographs in the four different background conditions displayed equivalent levels of light after digital alterations. However, this led to the question as to whether the artificial nature of digitally altered photo backgrounds may negatively impact sociability scores or online link clicking on pet profiles.

### Awareness of hypotheses

Participant awareness of study hypotheses did not influence link clicking on pet profiles or sociability scores of photographed shelter dogs. It was hypothesized that participant link clicking on pet profiles and sociability scores may be negatively impacted if photos appeared digitally altered. Past research shows that image manipulations involving splicing where photo elements differ from the original content can elicit feelings of deception by viewers in a context-dependent manner [23]; while image modifications for humans were considered generally acceptable for the purpose of photography, photo alterations for advertisements and journalism increased the perceived deception of viewers [23].

In relation to photo alterations for marketing strategies of shelter dogs, the results suggest that digitally altered backgrounds of online dog photographs may be acceptable in the context of sheltering. In fact, participants who were fully aware of the artificial nature of the photographs displayed increased link clicking behaviour, although this was not statistically significant. Participants who were definitely (score 5) and probably (score 4) aware of study hypotheses, and therefore image manipulation, had higher mean sociability scores compared to participants who were probably not (score = 2) or might or might not (score = 3) aware, although this was similarly not statistically significant. While previous studies have assessed the impact of digital enhancements of shelter photos (e.g. frames, text) on the speed of adoption of shelter dogs [12], no other study has specifically determined the effects of digital background alterations on viewer online interest and perceived sociability from photographed dogs.

While the results suggest that there is no positive impact of background types on sociability scores or link clicking behaviour, altering backgrounds of online photos of shelter dogs may be a potential tool for improving the speed of adoption by facilitating photo-taking of shelter dogs within foster homes. Foster families can provide online marketing materials of foster dogs without displaying their home environments. They can preserve their at-home privacy from the public by taking advantage of background modifying photo apps that are readily available online. However, further research is needed to directly assess the effectiveness of foster families utilizing tools to alter background types of photographed foster dogs.

### Morphological traits

Overall, link clicking behaviour and sociability scores were largely driven by dog ID suggesting that online viewer interest in online photos of shelter dogs is more influenced by the appearance of dogs rather than background type. This is congruent with past literature where the appearance of animals was listed as one of the primary reasons for adoption [10]. It is possible that the attention of participants was drawn to more salient photo traits relating to the physical features of dogs. As the appearance of dogs remained a significant driver impacting link clicking and sociability scores despite participants having a single encounter with photographed dogs, these results provide further support for the strength of morphological traits in the online interest of potential adopters. Workman and Hoffman found that salient morphological traits such as coat colour resulted in more clicks on the pet profiles of cats [9]. Using eye-tracker data, Isgate and Couchman found 92.7% of participants directed their attention to the facial features of photographed dogs the fastest and for the longest period of time [24]. Specifically, the fixation on facial features may explain the difference in link clicking behaviour and sociability scores between dog pairs within this study. A commonality between photographed dog pairs that differed significantly for link clicking (Liberty-Rogue; Liberty-Anakin) and sociability scores (Anakin-Liberty; Anakin-Phantom; Anakin-Rogue; Liberty-Rogue; Phantom-Rogue) were the variations in facial expressions. Open-mouthed differences in these photographs likely contribute to differential assessment of sociability and link clicking between dogs as past research shows that the mouth region of photographed dogs receives the most attention [24]. It is possible that the extension of the tongue in photographs was perceived favourably by participants when scoring sociability as dogs may appear to be “smiling.” Past research shows that individuals that have significant or minimal experience working with dogs can both recognize and correctly associate facial expression of photographed dogs to the situation evoking the emotion (e.g. ball to stimulate happy expression) [25]. However, further research is required to determine how interpretations of facial expressions of photographed shelter dogs by potential adopters relates to link clicking on pet profiles and sociability scores.

### Participant-related factors

Participant-related factors such as age and prior or current dog ownership experience did not predict link clicking nor sociability scores of photographed shelter dogs in varying background types. This was unexpected as we hypothesized that these participant-related factors would influence the online interest of dogs displayed in different background types. The results are in contrast to research which shows that the decision to adopt a dog is influenced by previous dog ownership or participant age-related factors [26,27]. Common owner-related reasons influencing the decision to adopt included the replacement of a prior dog or the need for companionship either for another dog, children, the family, or for adults [27]. The results also suggest that the participants’ location of residence impacted sociability scores but not link clicking. In particular, sociability scores of photographed shelter dogs from the Americas was higher than Europe. This indicates potential cultural differences that may shape the perception of personality in shelter dogs. It is also possible that these results are driven by different preferences and motivations to adopt that are community and shelter specific [12,28]; however, further research is needed.

### Conclusion

Overall, this study contributes to the growing body of literature on the online marketing of shelter dogs. The results of this study indicate that photo backgrounds of shelter dogs may not be the primary initial focus for online viewers compared to more salient photo traits such as the appearance of shelter dogs. However, photo manipulations of backgrounds do not detract from online viewing interest nor the perceived assessment of sociability; using digitally altered backgrounds as a tool for foster families to conceal their home environments has potential benefits which warrants further research.

## Acknowledgements

We thank Lexis Ly, Dr. David Fraser, and the Animal Welfare Program at the University of British Columbia for their support and comments for this study. In addition, we thank the R group for their advice on statistical approaches as well as the Haven Animal Care Shelter in Lubbock, Texas for providing the photographs of Liberty, Phantom, Anakin, and Rogue.

## Supporting information

**S1 Table. Summary of sociability scores**. Sociability scores between dog ID and background type: sample size (N), mean, standard deviation, median, interquartile range (IQR).

**S2 Virtual Experiment Questions**

**S3 R Script code**

**S4 Data set**

## References

1. Pet statistics. ASPCA [Internet]. 2018 [cited 2021 Apr 2]. Available from: https://www.aspca.org/animal-homelessness/shelter-intake-and-surrender/pet-statistics

2. Animal shelter statistics. Humane Canada [Internet]. 2019 [cited 2021 Apr 25]. Available from: https://humanecanada.ca/wp-content/uploads/2020/11/Humane_Canada_Animal_shelter_statistics_2019.pdf

3. Woodruff K, Smith DR. An estimate of the number of dogs in US shelters in 2015 and the factors affecting their fate. J Appl Anim Welf Sci. 2020 Jul 2; 23(3):302–14.

4. Coppola CL, Grandin T, Enns RM. Human interaction and cortisol: Can human contact reduce stress for shelter dogs? Physiol Behav. 2006 Mar 30;87(3):537–41.

5. Hennessy MB. Using hypothalamic–pituitary–adrenal measures for assessing and reducing the stress of dogs in shelters: A review. Appl Anim Behav Sci. 2013 Dec 1;149(1):1–12.

6. Newbury S, Blinn MK, Bushby PA, Cox CB, Dinnage JD, Griffin B, et al. Guidelines for standards of care in animal shelters. Association of Shelter Veterinarians. 2010:1–64.

7. Computer and internet use in the United States. The United States Census Bureau [Internet]. 2016 [cited 2021 Apr 19]. Available from: https://www.census.gov/library/publications/2018/acs/acs-39.html

8. Use of Internet services and technologies by age group and household income quartile. Government of Canada Statistics Canada [Internet]. 2019 [cited 2021 Apr 25]. Available from: https://www150.statcan.gc.ca/t1/tbl1/en/tv.action?pid=2210011301

9. Workman MK, Hoffman CL. An evaluation of the role the internet site Petfinder plays in cat adoptions. J Appl Anim Welf Sci. 2015 Oct 2;18(4):388–97.

10. Weiss E, Miller K, Mohan-Gibbons H, Vela C. Why did you choose this pet?: Adopters and pet selection preferences in five animal shelters in the United States. Animals. 2012 Jun; 2(2):144–59.

11. Lampe R, Witte TH. Speed of dog adoption: Impact of online photo traits. J Appl Anim Welf Sci. 2015 Oct 2;18(4):343–54.

12. Nakamura M, Dhand N, Wilson BJ, Starling MJ, McGreevy PD. Picture perfect pups: How do attributes of photographs of dogs in online rescue profiles affect adoption speed? Animals. 2020 Jan;10(1):152.

13. Thorn JM, Mitchell JT. Factors that influence adopters’ preference of shelter dogs. J Vet Behav. 2013;4(8):41–2.

14. Fratkin JL, Baker SC. The role of coat color and ear shape on the perception of personality in dogs. Anthrozoös. 2013 Mar 1;26(1):125–33.

15. Wright JC, Smith A, Daniel K, Adkins K. Dog breed stereotype and exposure to negative behavior: Effects on perceptions of adoptability. J Appl Anim Welf Sci. 2007 Jun 11;10(3):255–65.

16. Protopopova A, Mehrkam LR, Boggess MM, Wynne CDL. In-kennel behavior predicts length of stay in shelter dogs. PLoS One. 2014 Dec 31;9(12):e114319.

17. Protopopova A, Wynne CDL. Adopter-dog interactions at the shelter: Behavioral and contextual predictors of adoption. Appl Anim Behav Sci. 2014 Aug 1;157:109–16.

18. Hall RH, Hanna P. The impact of web page text-background colour combinations on readability, retention, aesthetics and behavioural intention. Behav Inf Technol. 2004 May 1;23(3):183–95.

19. Viswanathan PK, Swaminathan TN. Quantifying the Relative Importance of Key Drivers of Landing Page. Indian J Mark. 2017 Nov 1;47(11):24–35.

20. Nakamura M, Dhand NK, Starling MJ, McGreevy PD. Descriptive texts in dog profiles associated with length of stay via an online rescue network. Animals. 2019 Jul;9(7):464.

21. Protopopova A, Gilmour AJ, Weiss RH, Shen JY, Wynne CDL. The effects of social training and other factors on adoption success of shelter dogs. Appl Anim Behav Sci. 2012 Dec 15;142(1):61–8.

22. Pyzer C, Clarke L, Montrose VT. Effects of video footage versus photographs on perception of dog behavioral traits. J Appl Anim Welf Sci. 2017 Jan 2;20(1):42–51.

23. Conotter V, Dang-Nguyen D-T, Boato G, Menéndez M, Larson M. Assessing the impact of image manipulation on users’ perceptions of deception. Human Vision and Electronic Imaging XIX. 2014 Feb 25; 9014:90140Y.

24. Isgate S, Couchman JJ. What makes a dog adoptable? An eye-tracking Investigation. J Appl Anim Welf Sci. 2018 Jan 2;21(1):69–81.

25. Bloom T, Friedman H. Classifying dogs’ (Canis familiaris) facial expressions from photographs. Behav Processes. 2013 Jun 1;96:1–10.

26. Holland KE. Acquiring a pet dog: A review of factors affecting the decision-making of prospective dog owners. Animals. 2019 Apr;9(4):124.

27. Marston LC, Bennett PC, Coleman GJ. Adopting shelter dogs: Owner experiences of the first month post-adoption. Anthrozoös. 2005 Dec 1;18(4):358–78.

28. Brown WP, Davidson JP, Zuefle ME. Effects of phenotypic characteristics on the length of stay of dogs at two no kill animal shelters. J Appl Anim Welf Sci. 2013 Jan 1;16(1):2–18.

29. Gunter LM, Barber RT, Wynne CDL. A canine identity crisis: Genetic breed heritage testing of shelter dogs. PLoS One. 2018 Aug 23;13(8):e0202633.

30. Salman MD, New, Jr. JG, Scarlett JM, Kass PH, Ruch-Gallie R, Hetts S. Human and animal factors related to relinquishment of dogs and cats in 12 selected animal shelters in the United States. J Appl Anim Welf Sci. 1998 Jul 1;1(3):207–26.

31. Bonnardel N, Piolat A, Le Bigot L. The impact of colour on website appeal and users’ cognitive processes. Displays. 2011 Apr 1;32(2):69–80.

32. Broeder P, Snijder H. Colour in online advertising: Going for trust, which blue is a must? Mark-Inf Decis J. 2019 Jun 1;2(1):5–15.

33. Cyr D, Head M, Larios H. Colour appeal in website design within and across cultures: A multi-method evaluation. Int J Hum-Comput Stud. 2010 Jan 1;68(1):1–21.

34. Lee S, Rao V. Color and store choice in electronic commerce: the explanatory role of trust. J Electron Commer Res. 2010 May 1;11(2):110–26.

35. Gunter LM, Barber RT, Wynne CDL. What’s in a name? Effect of breed perceptions & labeling on attractiveness, adoptions & length of stay for pit-bull-type dogs. PLoS One. 2016 Mar 23;11(3):e0146857.

36. R Core Team. R: A language and environment for statistical computing [Internet]. 2016 [cited 2021 Jun 1]. Available from: https://www.r-project.org/

